# Maternal immunization with distinct influenza vaccine platforms elicits unique antibody profiles that impact the protection of offspring

**DOI:** 10.1101/2023.10.30.564827

**Authors:** Ana Vazquez-Pagan, Ericka Kirkpatrick Roubidoux, Sean Cherry, Brandi Livingston, Theresa Bub, Lauren Lazure, Bridgett Sharp, Thomas Confer, Pamela H. Brigleb, Rebekah Honce, Kendall T. Whitt, Melissa Johnson, Victoria Meliopoulos, Stacey Schultz-Cherry

## Abstract

Pregnant women and infants are considered high-risk groups for increased influenza disease severity. While influenza virus vaccines are recommended during pregnancy, infants cannot be vaccinated until at least six months of age. Passive transfer of maternal antibodies (matAbs) becomes vital for the infant’s protection. Here, we employed an ultrasound-based timed-pregnancy murine model and examined matAb responses to distinct influenza vaccine platforms and influenza A virus (IAV) infection in dams and their offspring. We demonstrate vaccinating dams with a live-attenuated influenza virus (LAIV) vaccine or recombinant hemagglutinin (rHA) proteins administered with adjuvant resulted in enhanced and long-lasting immunity and protection from influenza in offspring. In contrast, a trivalent split-inactivated vaccine (TIV) afforded limited protection in our model. By cross-fostering pups, we show the timing of antibody transfer from vaccinated dams to their offspring (prenatal versus postnatal) can shape the antibody profile depending on the vaccine platform. Our studies provide information on how distinct influenza vaccines lead to immunogenicity and efficacy during pregnancy, impact the protection of their offspring, and detail roles for IgG1 and IgG2c in the development of vaccine administration during pregnancy that stimulate and measure expression of both antibody subclasses.

## Introduction

For over a century, pregnancy has been acknowledged as a risk factor for severe complications from various infectious agents including influenza virus.^1,2^ Epidemiological and empirical studies have shown influenza virus infection during pregnancy can result in increased disease severity, including a five-fold increase in cardiopulmonary complications, as compared to a non-pregnant woman.^3^ Mortality rates were also highest among pregnant women (as high as 45%) during major influenza pandemics (1918, 1957, 1968, and 2009).^3–6^ This risk is not limited only to pregnant women as infection-related complications also extend to their offspring. These complications include an increased risk of miscarriage, stillbirth, neonatal death, preterm birth, and low birth weight.^5–14^

Given the plethora of adverse outcomes that can result from gestational influenza infection, prevention through vaccination is the best control strategy and has been heavily advocated for by the World Health Organization (WHO). Several different influenza virus vaccines are administered annually as the standard of care (SOC). These include a trivalent or quadrivalent inactivated vaccine (TIV/QIV), live-attenuated influenza vaccine (LAIV), and recombinant hemagglutinin (rHA) protein. It is recommended for pregnant women to receive an influenza vaccine in the form of TIV or QIV anytime during pregnancy, with an emphasis on vaccination early in an influenza season.^15^ An additional advantage of maternal immunization is it can protect newborns as they are not equipped with a mature immune system that can protect them from infections.^16,17^ The transfer of maternal antibodies (matAbs) during pregnancy or breastfeeding becomes crucial for the infant’s protection against infections in early life.^18–20^LLPrevious work has evaluated whether influenza A virus (IAV) infection or vaccination with a split-inactivated vaccine results in prolonged protection in the offspring.^21–23^ However, there is a lack of studies evaluating other influenza vaccine delivery platforms administered during pregnancy and how these may influence passive immunity.

To fill this knowledge gap, we established a novel ultrasound-based mouse pregnancy model and used it to evaluate the efficacy and immunogenicity of distinct influenza vaccine platforms administered during pregnancy and how these may influence the response to influenza infection in the offspring. Our studies demonstrated that vaccinating dams with LAIV or rHA proteins administered with adjuvant resulted in enhanced and long-lasting immunity and protection in dams and offspring, similar to what is observed in response to infection. In comparison, TIV afforded limited protection. Cross-fostering showed that the timing of antibody transfer (prenatal or postnatal) and the antibody profiles from the vaccinated dams to pups differ depending on the vaccine platform. These studies aided in elucidating the underlying mechanisms of matAb-mediated protection for distinct influenza vaccine platforms yet could also extend to other respiratory viruses of similar pathology and advance vaccine development pipelines specific to maternal immunization.

## Results

### Vaccination does not lead to adverse outcomes in pregnant mice

Information on the efficacy and immunogenicity of influenza vaccination administered during pregnancy is limited to inactivated vaccines and is primarily focused on protection in the dams.^25^ To investigate how distinct influenza vaccine platforms impact dams and offspring, we confirmed pregnancy by ultrasound. Dams were primed on embryonic day (E) 8.0 with the standard of care (SOC) TIV, or a cocktail of recombinant hemagglutinin proteins (rHAs) mixed 1:1 with Addavax administered intramuscularly and boosted 3 weeks later (**Figure 1A**). In addition, two subsets of pregnant dams were inoculated intranasally with a sublethal dose of A/California/04/2009 H1N1 virus (IAV) or with cold-adapted A/California/04/2009 H1N1 live attenuated virus (LAIV) at E8.0. Non-vaccinated pregnant mice served as controls.

**Figure 1.**
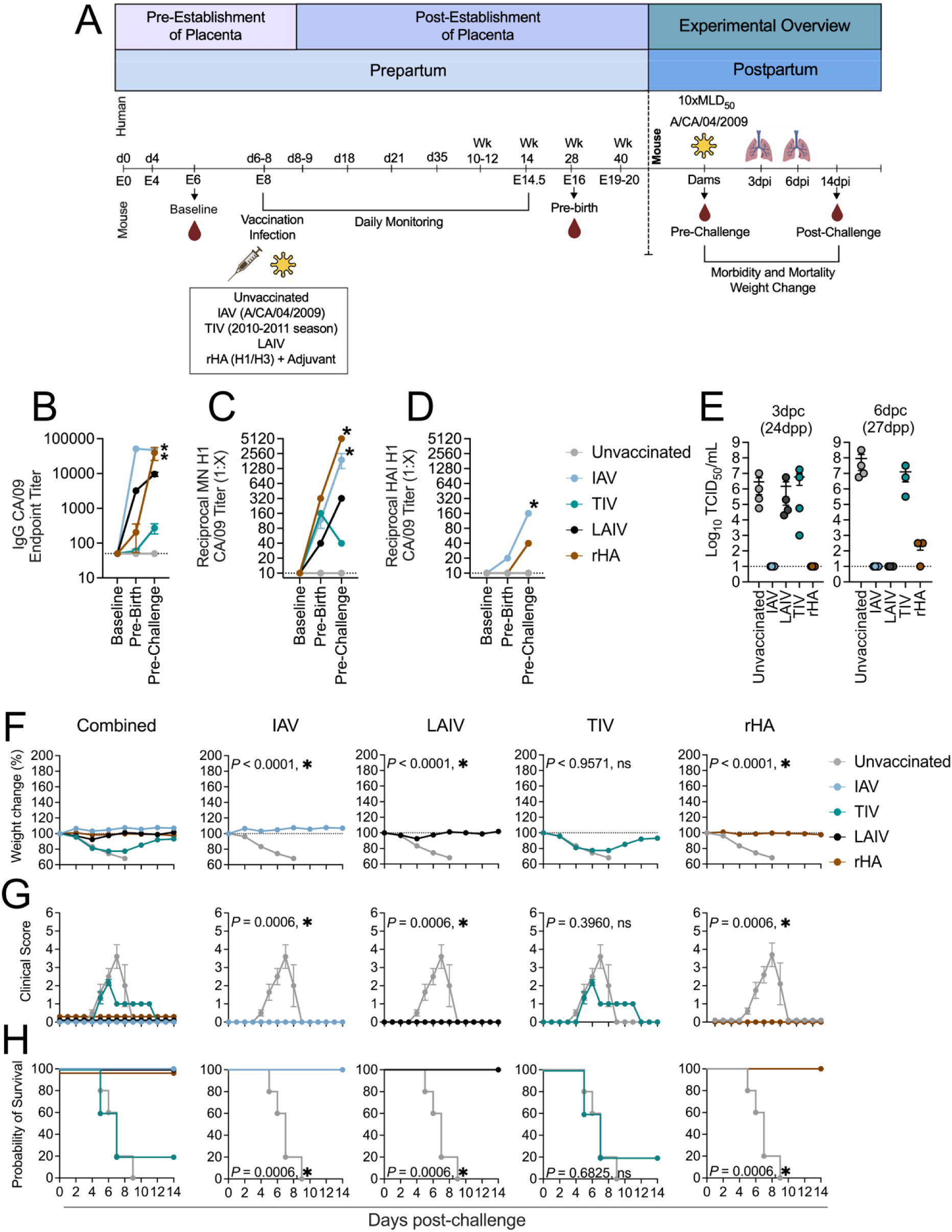
Vaccination platforms differ in efficacy and immunogenicity. (A) Vaccination scheme and sample collection timeline. Antibody levels in longitudinally collected sera were quantified via (B) viral-specific ELISA (IAV p<0.0001, TIV p>0.9999, LAIV p=0.8346, rHA p=0.0041), (C) microneutralization (MN) assay (IAV p=0.0336, TIV p=0.3805, LAIV p=0.3555, rHA p<0.0001), and (D) hemagglutination inhibition (HAI) assays (IAV p=0.0336, TIV p=0.3805, LAIV p=0.3555, rHA p<0.0001) compared via two-way ANOVA (E) Lung viral load at 3 days post-challenge (dpc), or day 24 days post-partum (dpp) and day 6 dpc, or 27 dpp. Grey dotted lines indicate a lower limit of detection. (F-H) Morbidity and mortality in vaccinated or infected pregnant mice compared to unvaccinated mice with n=10 mice/group. (F) Weight curves post-challenge with statistical comparisons made via mixed-effects model. Grey dotted line indicates a 0% deviation from weight at baseline. (G) Clinical scores post-challenge with statistical comparisons made using a two-way ANOVA. (H) Survival post-challenge with statistical comparisons using Mantel-Cox log-rank analysis. Data represent three independent experiments and are graphed as means ± standard error and in (H) as surviving proportions with censored or event animals indicated by their respective symbols. Statistical comparison tests are between unvaccinated animals and indicated vaccination or infection group (n=15 mice/group) in (B-D), (n=3-5 mice/group) in (E), and (n=10 mice/group) in (F-H).

No adverse pregnancy outcomes (i.e., substantial body weight fluctuations or adverse fetal outcomes) were observed in the TIV and rHA-vaccinated groups. Dams infected with a sub-lethal dose of IAV failed to gain as much weight as non-infected dams, typically, between 10-20% less, and had minor clinical signs of infection (**Supplementary Figure 1A-B**). Similar results were observed in IAV-inoculated non-pregnant female mice (**Supplementary Figure 1C-D**). In contrast, LAIV-vaccinated mice exhibited no clinical signs of infection and pregnant mice gained weight like unvaccinated controls (**Supplementary Figure 1**) illustrating that infection poses a higher risk to dam health during pregnancy than vaccination, even at a low inoculation dose.

### Immunogenicity and efficacy differ among influenza vaccine platforms in immunized dams

To assess influenza-specific antibody responses for each vaccine platform, viral-specific IgG ELISA, microneutralization (MN), and hemagglutination inhibition (HAI) titers were quantified at baseline, before delivery (pre-birth), and pre-challenge, which corresponds to five weeks post-vaccination, or 21 days postpartum (dpp) (**Figure 1B-D, Supplementary Table 1**). Pregnant mice vaccinated with LAIV, rHA, or infected with IAV, had the highest viral-specific ELISA IgG, MN, and HAI titers pre-birth and challenge compared to TIV-vaccinated and unvaccinated dams (**Figure 1B-D, Supplementary Table 1**). At the pre-boost time point, viral-specific ELISA and MN IgG titers were lowest in the TIV-vaccinated group but did increase by the pre-challenge time point. However, antibody titers in this group never reached similar levels as sub-lethally infected, LAIV, or rHA-vaccinated dams but were higher than unvaccinated controls (**Figure 1B-C, Supplementary Table 1**). Of note, while viral-specific ELISA and MN IgG titers were detected in all groups, albeit at distinct levels, only IAV-infected and rHA-vaccinated dams had detectable HAI titers (**Figure 1D, Supplementary Table 1**). These data illustrate the variable magnitude and quality of antibody responses elicited by different influenza vaccine platforms or infection during pregnancy.

To determine which influenza vaccine platform would provide the best protection, dams were challenged with 10x mouse lethal dose-50 (MLD_50_) A/California/04/2009 (A/CA/04/2009) H1N1 virus as indicated in **Figure 1A** and monitored for morbidity and mortality for fourteen days (**Figure 1E-H**). Unlike the unvaccinated group, LAIV and rHA-vaccinated dams had little to no weight loss, statistically reduced signs of infection and 100% survival (**Figure 1F-H**) following the viral challenge. This was also found in dams sub-lethally infected with IAV who exhibited 100% survival and reduced morbidity (**Figure 1F-H**) upon secondary challenge. In contrast, TIV administered early during pregnancy failed to fully protect the post-partum dams. The TIV-vaccinated dams lost around 20% of their body weight (**Figure 1F**) and had clinical scores comparable to unvaccinated mice (**Figure 1G**). Yet, TIV-vaccinated dams were still better protected than unvaccinated dams, however, this was not statistically significant (**Figure 1H**).

To better understand why the survival outcomes varied between vaccine platforms, we monitored lung viral titers post-challenge (**Figure 1E, Supplementary Table 1**). At 3 days post-challenge (dpc), reduced viral titers were only observed in IAV-infected and rHA-vaccinated dams, which aligned well with high HAI and neutralizing antibody titers (**Figures 1C-D, Supplementary Table 1**). In contrast, TIV and LAIV-vaccinated dams exhibited high lung viral titers at 3 dpc similar to unvaccinated controls (**Figure 1E, Supplementary Table 1**). However, by 6 dpc TIV-vaccinated dams were the only group to exhibit high viral titers, albeit still lower than unvaccinated controls (**Figure 1E, Supplementary Table 1**), suggesting that neutralizing antibodies play a vital role in viral clearance, and ultimately survival to challenge.

### TIV immunogenicity improves in non-pregnant female mice

Pregnant women’s altered responses to infectious diseases are well-established,^2^, where it is known that a complex interplay between sex hormones and immunity drives different responses between non-pregnant and pregnant females.^23–26^ To determine if the differential vaccine responses were due to the pregnant status of the dams when immunized, age-matched non-pregnant C57BL/6J female mice were subjected to the same vaccination and challenge protocols (**Figure 2A**). Non-pregnant female mice immunized with rHA, or inoculated with LAIV, or IAV, exhibited higher viral-specific ELISA IgG and MN titers compared to unvaccinated or TIV-vaccinated non-pregnant female mice (**Figures 2B-D**). However, TIV-vaccinated non-pregnant mice did exhibit higher viral-specific ELISA and MN IgG titers compared to mice vaccinated while pregnant (**Figures 1B-D**, **Figures 2B-D**). Significant weight loss or presence of clinical scores after challenge for IAV-infected or rHA- and LAIV-vaccinated non-pregnant females was not observed (**Figures 2F-H**). In contrast, TIV-vaccinated female mice experienced infection-induced weight loss and higher clinical scores compared to the other groups and were not significantly different than that of unvaccinated mice (**Figure 2F-2G**).

**Figure 2.**
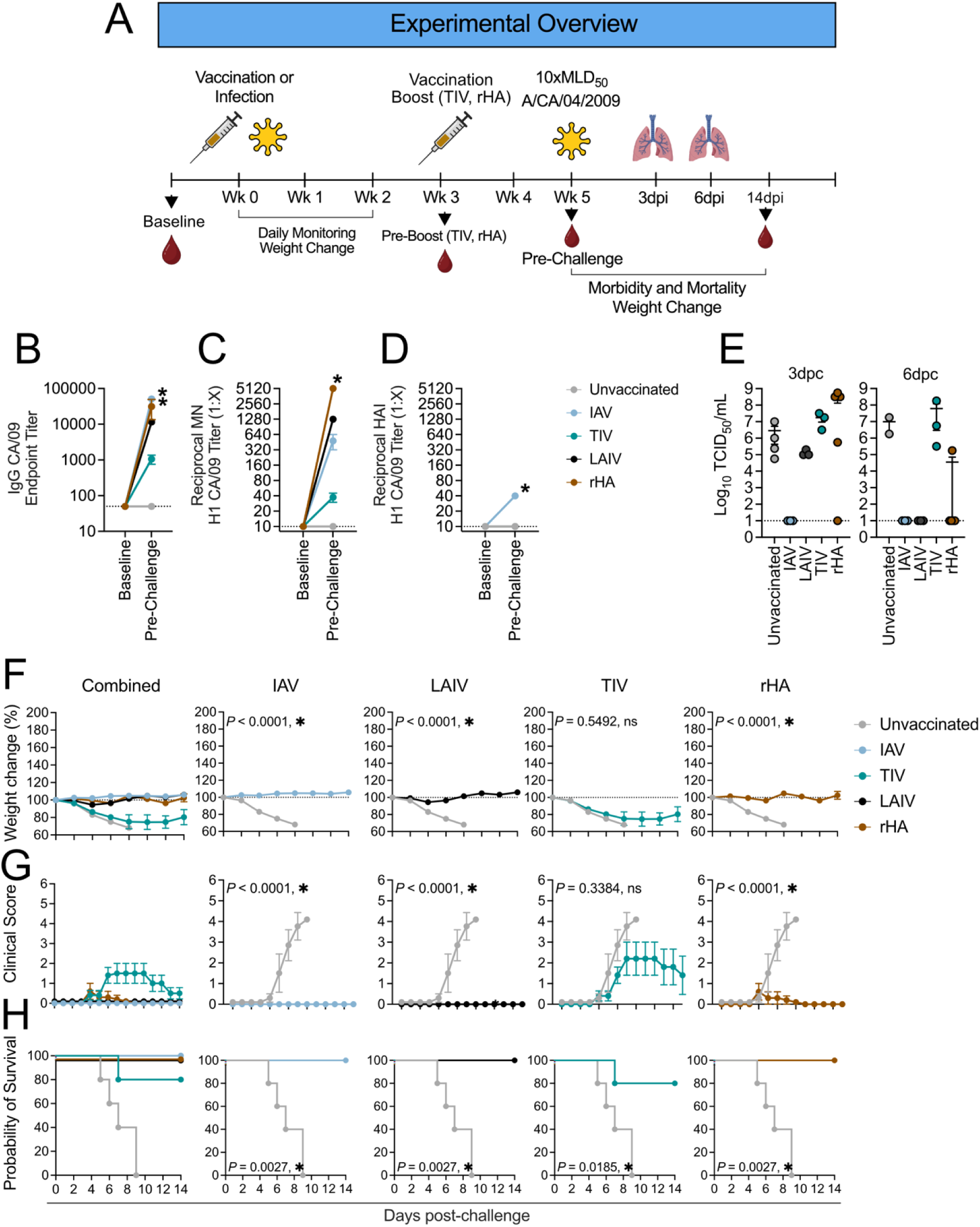
Vaccine efficacy and immunogenicity in non-pregnant female mice. (A) Vaccination scheme. Longitudinal antibody levels in sera were quantified via (B) viral-specific ELISA (IAV p<0.0001, TIV p=0.7698, LAIV p=0.9084, rHA p=0.0619), (C) MN assay (IAV p=0.9974, TIV p=0.9173, LAIV p=0.7874, rHA p=0.0020), and (D) hemagglutination inhibition (HAI) assay (IAV p=0.0061, TIV p>0.9999, LAIV p>0.9999, rHA p>0.9999) compared via two-way ANOVA. (E) Lung viral load at 3 days post-challenge (dpc), or day 6 dpc. Grey dotted lines indicate a lower limit of detection. (F-H) Morbidity and mortality in vaccinated or infected pregnant mice compared to unvaccinated mice with n=10 mice/group. (F) Weight curves post-challenge with statistical comparisons made via mixed-effects model. Grey dotted line indicates a 0% deviation from weight at baseline. (G) Clinical scores post-challenge with statistical comparisons made using a two-way ANOVA. (H) Survival post-challenge with statistical comparisons using Mantel-Cox log-rank analysis. Data represent three independent experiments and are graphed as means ± standard error and in (H) as surviving proportions with censored or event animals indicated by their respective symbols. Statistical comparison tests are between unvaccinated animals and indicated vaccination or infection group (n=15 mice/group) in (B-D), (n=3-5 mice/group) in (E), and (n=5-8 mice/group) in (F-H).

A sub-lethal homologous IAV infection or vaccination with LAIV and rHA significantly reduced mortality (**Figure 2H**) upon challenge. While TIV vaccination resulted in reduced protection from a lethal influenza challenge as compared to other influenza vaccine platforms, mice were still significantly more protected against mortality compared to unvaccinated controls (**Figures 2F-H**). When evaluating viral load in the lungs of non-pregnant mice, the trend for IAV-infected and LAIV-vaccinated female mice mirrors that of mice immunized while pregnant (**Figure 1E**, **Figure 2E**). In contrast, non-pregnant female mice vaccinated with rHA had high viral titers at 3 dpc, a clear disparity from dams immunized while pregnant (**Figure 1E**, **Figure 2E**). However, by 6 dpc, there is a trend toward viral clearance in this group (**Figure 2E**). Overall, TIV vaccination resulted in an 80% probability of survival for non-pregnant female mice compared to 20% survival in post-partum dams vaccinated during pregnancy. This suggests that there are pregnancy-induced alterations in the elicitation of a humoral response that are specific to the split-inactivated influenza vaccine platform. This is further illustrated by the blunted seroconversion during pregnancy and poor survival upon challenge post-partum (**Figures 1B-D, 1H**).

### Maternal vaccination-induced protection of 3-week-of-age offspring is dependent on the vaccine platform

In humans, infants younger than six months of age cannot receive an influenza vaccine, rendering this group highly susceptible to infection.^14^ However, transplacental transfer of matAbs during gestation or post-birth through breast milk can protect offspring from early life exposures during this vaccination gap.^18–20^ Various studies have evaluated the ability of matAbs to boost the offspring’s humoral response to immunization or the role of matAbs in the interference of response to vaccination.^22,23,27,28^ However, few studies to date have evaluated the efficacy across influenza vaccine platforms nor their ability to confer long-term matAb-mediated protection to offspring.^28,29^

To better understand the magnitude and kinetics of matAbs elicited by distinct influenza vaccine platforms, we evaluated viral-specific ELISA IgG, MN, and HAI titers in weanlings born to vaccinated, infected, and unvaccinated dams (**Figures 3A-3D**). At weaning, we observed approximately 10-fold lower ELISA titers in three-week-old pups compared to dams (**Figure 3B, Supplementary Table 2**). Neutralizing antibodies were also detected at lower levels in the three-week-old pups than in dams, except for offspring born to rHA dams (**Figure 3C, Supplementary Table 2**). Three-week-old pups born to IAV-infected dams had the highest viral-specific ELISA IgG titers compared to the other groups (**Figure 3B, Supplementary Table 2**) and were the only group with detectable HAI titers (**Figure 3C, Supplementary Table 2**). In contrast, pups born to rHA-vaccinated dams had the highest neutralizing antibody titers as compared to other platforms (**Figure 3D, Supplementary Table 2**), reflecting the same trend observed in the dams (**Figure 1D, Supplementary Table 1**). Overall, the magnitude and neutralizing activity of antibodies in three-week-old pups varied by vaccine platform and mirrored the responses observed in the dams suggesting that different influenza vaccine platforms may elicit distinct antibody profiles and variable antibody transfer patterns in our model.

**Figure 3.**
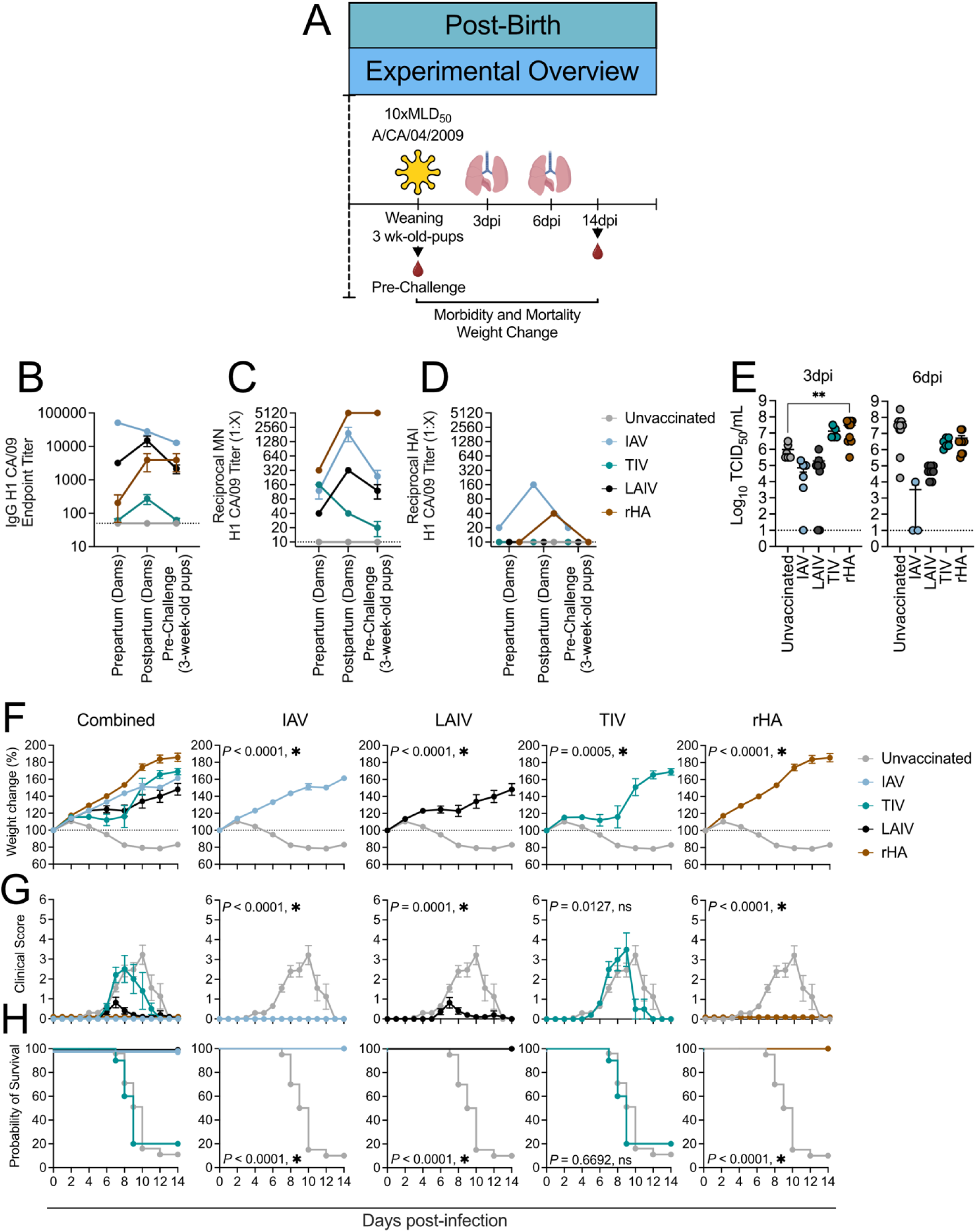
The vaccine platform impacts the efficacy and immunogenicity of maternal antibodies transferred to weanlings. (A) Timeline of weaning, infection, and harvests in three-week-old offspring born to vaccinated or infected dams. Longitudinal antibody levels in sera were quantified via (B) viral-specific ELISA (IAV p<0.0001, TIV p=0.7698, LAIV p=0.9084, rHA p=0.0619), (C) MN assay (IAV p=0.9974, TIV p=0.9173, LAIV p=0.7874, rHA p=0.0020), and (D) hemagglutination inhibition (HAI) assay (IAV p=0.0061, TIV p>0.9999, LAIV p>0.9999, rHA p>0.9999) compared via two-way ANOVA. (E) Lung viral load at 3 days post-infection (dpi; IAV p=0.9999, TIV p=0.3948, LAIV p>0.9999, rHA p=0.0009), or 6 dpi (IAV p=0.2950, TIV p=0.1574, LAIV p=0.0826, rHA p=0.1003). Grey dotted lines indicate a lower limit of detection. (F-H) Morbidity and mortality in pups born to vaccinated or infected dams as compared to pups born to unvaccinated mice with n=10 mice/group. (F) Weight curves post-infection with statistical comparisons made via a mixed-effects model. Grey dotted line indicates a 0% deviation from weight at baseline. (G) Clinical scores post-infection with statistical comparisons made using a two-way ANOVA. (H) Survival post-infection with statistical comparisons using Mantel-Cox log-rank analysis. Data represent three independent experiments and are graphed as means ± standard error and in (H) as surviving proportions with censored or event animals indicated by their respective symbols. Statistical comparison tests are between unvaccinated animals and indicated vaccination or infection group (n=15 mice/group) in (B-D), (n=3-5 mice/group) in (E), and (n=10 mice/group) in (F-H).

To assess efficacy, three-week-old pups were challenged with the 10xMLD_50_ A/CA/04/2009 H1N1 virus (**Figures 3F-H**). Pups born to IAV-infected, rHA-, and LAIV-vaccinated dams had no body weight loss and no clinical scores leading to 100% survival as compared to offspring from unvaccinated controls (**Figures 3F-G**). In contrast, offspring born to TIV-vaccinated dams had arrested growth, high clinical scores, and a 20% probability of survival. These findings paired well with the levels of anti-influenza IgG and neutralizing antibodies present at the time of infection (**Figures 3B-D**). To examine the efficacy of matAbs in blocking viral replication, we quantified whole-lung viral titers at 3- and 6-dpi. Offspring born to IAV-infected and LAIV-vaccinated dams had reduced lung viral titers at 3 dpi with clearance in IAV and reduced levels in LAIV-vaccinated dams by 6 dpi, which agrees with the complete survival observed in these groups (**Figures 3E-H, Supplementary Table 2**). In contrast, offspring born to rHA- and TIV-vaccinated dams had the highest viral load at 3 dpc with limited reduction at 6 dpi (**Figure 3E**).

While offspring born to TIV- and rHA-vaccinated dams still had high viral load as compared to the other groups, titers were overall lower than pups born to unvaccinated dams (**Figure 3E**). Increased viral burden in the lungs of offspring born to TIV-vaccinated dams aligned with the absence of matAbs before infection (**Figure 3B-D**) and the poor survival observed in this group (**Figure 3H**). Despite having high lung viral titers at 3 and 6 dpi, the offspring of rHA-vaccinated dams were protected, suggesting matAbs are playing a role in controlling the infection through a non-neutralizing mechanism (**Figure 3H**). Overall, these studies demonstrate that the type of vaccine used during pregnancy can significantly impact the efficacy and immunogenicity of the response to a lethal influenza challenge in weanlings.

### Maternally transferred antibodies persist until eight weeks of age and protect offspring born to rHA-vaccinated dams

To assess the longevity of the antibody response, the offspring of infected and vaccinated dams were monitored for 5 additional weeks post-birth and then infected as described (**Figure 4A**). The offspring were weighed weekly, and we did not observe any significant differences between groups in terms of pup size as they aged (**Figure 4B**). Viral-specific ELISA IgG and MN titers remained detectable and above baseline until eight weeks of age in pups born to IAV-infected, rHA-, and LAIV-vaccinated dams. However, in pups born to TIV-vaccinated dams, viral-specific ELISA IgG and MN titers waned by six weeks of age (**Figures 4C-E**), a finding recapitulating previous studies.^27^

**Figure 4.**
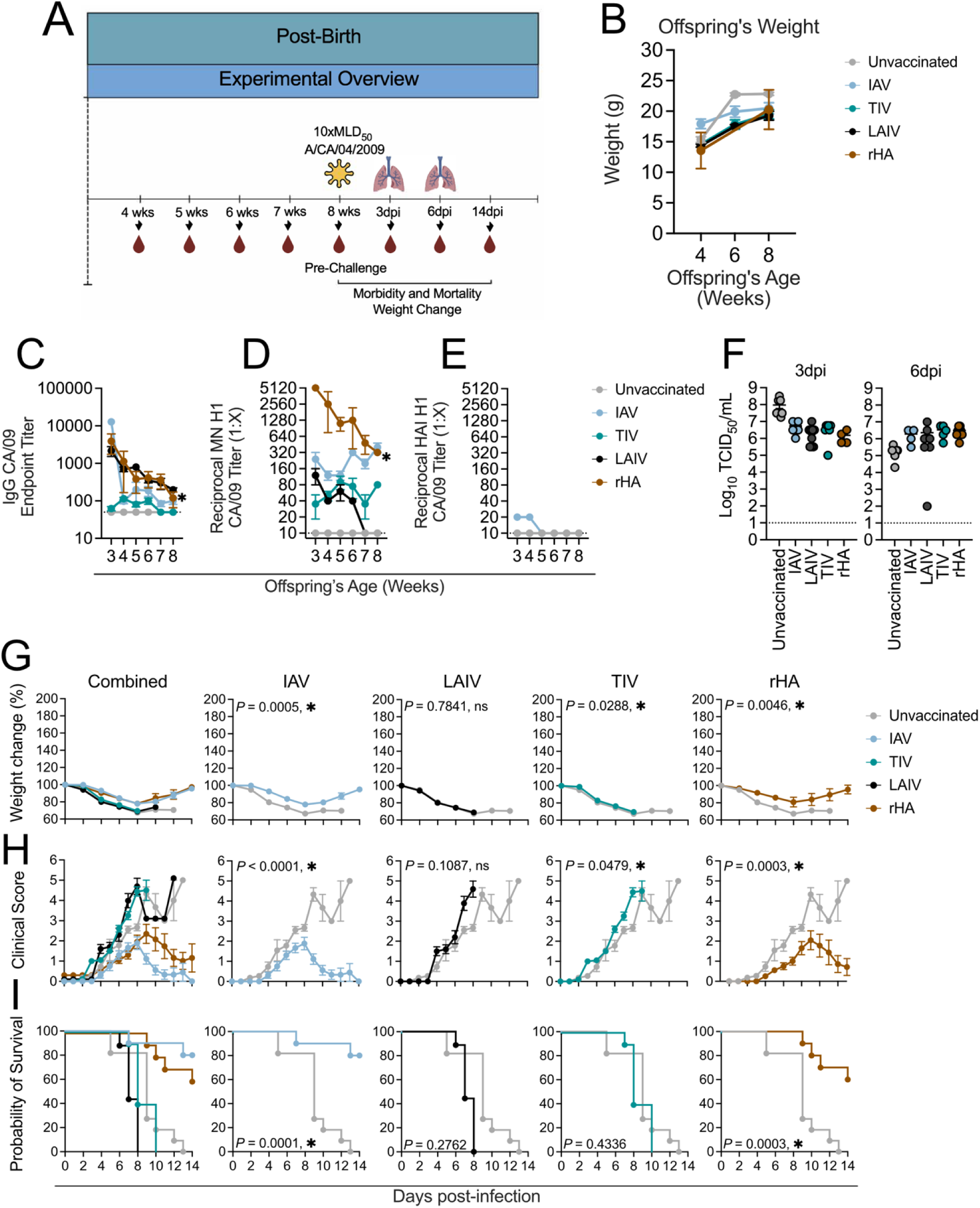
Longitudinal antibody responses over time in offspring born to vaccinated dams. (A) Experimental overview and challenge timeline. (B) Weekly weights of offspring born to vaccinated or infected dams. Longitudinal antibody levels in sera were quantified via (C) viral-specific ELISA (IAV p<0.0001, TIV p=0.7652, LAIV p=0.9996, rHA p=0.3580), (D) MN assay (IAV p=0.5392, TIV p=0.9999, LAIV p=0.9997, rHA p=0.0020), and (E) hemagglutination inhibition (HAI) assay (IAV p=0.0061, TIV p>0.9999, LAIV p>0.9999, rHA p<0.0001) compared via two-way ANOVA. (F) Lung viral load at 3 days post-infection (dpi; IAV p=0.0104, TIV p=0.0081, LAIV p=0.0078, rHA p=0.0164), or 6 dpi (IAV p=0.7557, TIV p=0.1356, LAIV p=0.3826, rHA p=0.0843). Grey dotted lines indicate a lower limit of detection. (G-I) Morbidity and mortality in pups born to vaccinated or infected dams as compared to pups born to unvaccinated mice with n=10 mice/group. (G) Weight curves post-infection with statistical comparisons made via a mixed-effects model. Grey dotted line indicates a 0% deviation from weight at baseline. (H) Clinical scores post-infection with statistical comparisons made using a two-way ANOVA. (I) Survival post-infection with statistical comparisons using Mantel-Cox log-rank analysis. Data represent two independent experiments and are graphed as means ± standard error and in (I) as surviving proportions with censored or event animals indicated by their respective symbols. Statistical comparison tests are between pups born to unvaccinated dams and the indicated group of pups born to vaccinated dams (n=15 mice/group) in (B-E), (n=3-5 mice/group) in (F), and (n=10 mice/group) in (G-I).

To assess efficacy, we challenged eight-week-old pups with 10xMLD_50_ A/CA/04/2009 H1N1 virus (**Figures 4G-I**). Offspring born to IAV-infected dams exhibited mild clinical scores and 80% survival (**Figure 4G-I**), which is less than what was observed at 3 weeks of age. Offspring born to TIV-vaccinated dams experienced morbidity and mortality like pups born to unvaccinated dams (**Figures 4G-I**). Similar results were obtained in pups born to LAIV-vaccinated dams despite detectable viral-specific IgG ELISA titers (**Figures 4G-I**) but could be explained by a loss of neutralizing antibodies at 7 weeks. In contrast, offspring born to rHA-vaccinated dams were protected from infection although survival reduced from 100% at 3 weeks of age (**Figure 3H**) to 60%; significantly higher than offspring born to unvaccinated controls (**Figure 4H**). No reduction in lung viral load was observed in any of the groups at 3 and 6 dpi, suggesting that antibody levels and functionality were not sufficient to control viral replication (**Figure 4F**), and ultimately led to reduced survival. Altogether, these findings highlight that the platform used to vaccinate pregnant dams can influence the efficacy, immunogenicity, and longevity of the protective responses in the offspring.

### Cross-fostering highlights that the matAbs transferred prenatally mediate protection in 3-week-of-age offspring after exposure to a live virus

We questioned whether matAbs responsible for mediating protection in the offspring were transferred prenatally or postnatally via suckling (**Figure 5, Supplementary Table 2**). To test, pups from control unvaccinated, IAV-infected, or LAIV-, TIV- and rHA-vaccinated dams were cross-fostered at birth as described (**Figure 5A**). After three weeks (i.e., at the time of weaning), offspring born to infected or vaccinated dams nursed by unvaccinated controls (prenatal transfer) and naïve offspring nursed by infected or vaccinated dams (postnatal transfer) were challenged with 10xMLD_50_ A/CA/04/2009 H1N1 virus and monitored for morbidity and mortality (**Figure 5F-5H**).

**Figure 5.**
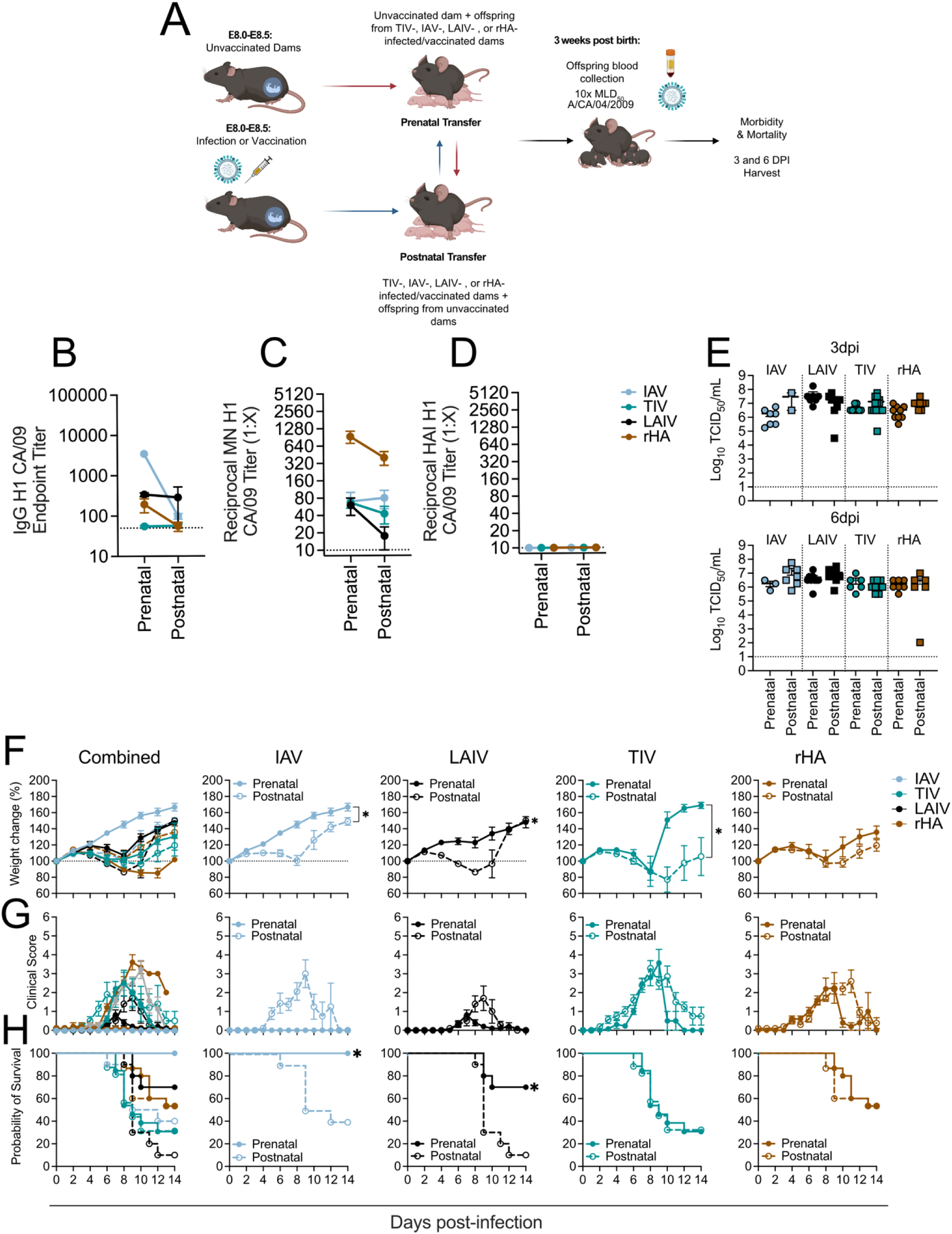
Cross-fostering of litters reveals matAb transfer is more efficient at the prenatal stage. (A) Schematic of cross-fostering experimental design. Antibody levels in sera were quantified via (B) viral-specific ELISA, (C) microneutralization (MN) assay and (D) hemagglutination inhibition (HAI) assay. (E) Lung viral load at 3- and 6-days post-infection (dpi). Grey dotted lines indicate a lower limit of detection. (F-H) Morbidity and mortality in pups with prenatal matAb transfer as compared to postnatal matAb transfer with n=5-10 mice/group. (F) Weight curves post-infection with statistical comparisons made via a mixed-effects model. Grey dotted line indicates a 0% deviation from weight at baseline. (G) Clinical scores post-infection with statistical comparisons made using a two-way ANOVA. (H) Survival post-infection with statistical comparisons made using Mantel-Cox log-rank analysis. Data in (B-H) are representative of two independent experiments and are graphed as means ± standard error and in (H) as surviving proportions with censored or event animals indicated by their respective symbols. Statistical comparisons tests are between pups that had matAb prenatal transfer and pups that received matAb postnatal (n=10 mice/group) in (B-D), (n=3-5 mice/group) in (E), and (n=8-10 mice/group) in (F-H).

Pups born to IAV-infected dams but nursed by unvaccinated dams (prenatal transfer) were protected from infection exhibiting the highest viral-specific IgG ELISA titers (**Figure 5B, Supplementary Table 2**), no morbidity (weight loss and clinical scores) and 100% survival (**Figures 5F-5H**). In contrast, offspring born to non-vaccinated mice and nursed by IAV-infected mice (postnatal transfer), had significantly increased morbidity (**Figures 5F-5G**) resulting in only 40% of mice surviving influenza infection (**Figure 5H**). While offspring born to LAIV-vaccinated dams (prenatal transfer) experienced minimal body weight loss upon infection (**Figure 5F**), they did experience increased morbidity, albeit not significant (**Figure 5G**), which resulted in less than 80% survival in this group (**Figure 5H**). Offspring born to unvaccinated mice that were nursed by LAIV-vaccinated dams (postnatal transfer), exhibited significant infection-related weight loss (**Figure 5F**), increased morbidity (**Figure 5G**), and significantly increased mortality resulting in less than 20% of offspring surviving post-infection as compared to pups born to LAIV-vaccinated dams (**Figures 5H**). Overall, these findings suggest protection in offspring born to IAV-infected and LAIV-vaccinated dams is primarily mediated upon matAbs transferred prenatally, rather than via suckling.

Pups from unvaccinated dams nursed by TIV dams (postnatal transfer) experience significantly increased weight loss as compared to pups who receive the prenatal transfer of matAbs (**Figure 5F**). However, the survival of both groups is comparable and not statistically significant from each other (**Figure 5H**). Offspring nursed by rHA-vaccinated dams (postnatal transfer) exhibited similar mortality as those who received a prenatal transfer of matAbs culminating in less than 60% survival for both groups despite having the highest presence of neutralizing antibodies (**Figure 5C, 5F-H, Supplementary Table 2**). Upon evaluation of viral burden in whole lungs at 3 and 6 dpi, no significant differences between any of the groups were observed (**Figure 5E)**, indicating that protection from infection with a lethal influenza dose does not appear to be mediated by rapid viral clearance, but instead by the ability of matAbs to control clinical disease. These data suggest that prenatal transfer of matAbs is responsible for the protection observed in offspring born to LAIV-vaccinated or IAV-infected dams, most likely via the presence of higher ELISA and MN titers transferred during this time (**Figures 5B-5C, Supplementary Table 2**). However, when dams are vaccinated with TIV or rHA, neither prenatal nor postnatal matAb transfer alone is sufficient to achieve complete protection, or 100% survival, in their offspring. Our data suggest that both prenatal and postnatal matAb transfer is needed in these groups to mediate the 100% protection observed in the 3-week-old offspring born to rHA-vaccinated dams (**Figures 5F-5H**, **Figures 3F-3H, Supplementary Table 2**).

### Maternally transferred antibodies can differ in antibody subclass profiles and vary by influenza vaccine platform

Previous studies have shown that in C57BL/6J mice, IgG2c is the subclass that is produced in the highest titers and affords the greatest level of protection against a secondary infection with IAV after vaccination.^30–35^ We reasoned that differences in the protective effect of the prenatal or postnatal matAbs transferred to offspring could be attributed to variations in the abundance of distinct IgG subclass antibodies. To test, we measured viral-specific IgG titers for subclasses IgG1, IgG2b, and IgG2c in the serum of pups prior to challenge with a lethal influenza dose (**Figure 5A**, **Figure 6, Supplementary Table 3**). Offspring born to IAV-infected and LAIV-vaccinated dams and nursed by unvaccinated mice (prenatal transfer) had the highest IgG1, IgG2b, and IgG2c viral-specific ELISA titers, with the IAV group having a greater than 5-fold increase for all IgG subclasses, as compared to LAIV (**Figures 6A-B, Supplementary Table 3**). The discrepancy in the magnitude of the antibody response between IAV and LAIV groups aligns well with the observed differences in survival for offspring from these infected or vaccinated dams (**Figure 5F**).

**Figure 6.**
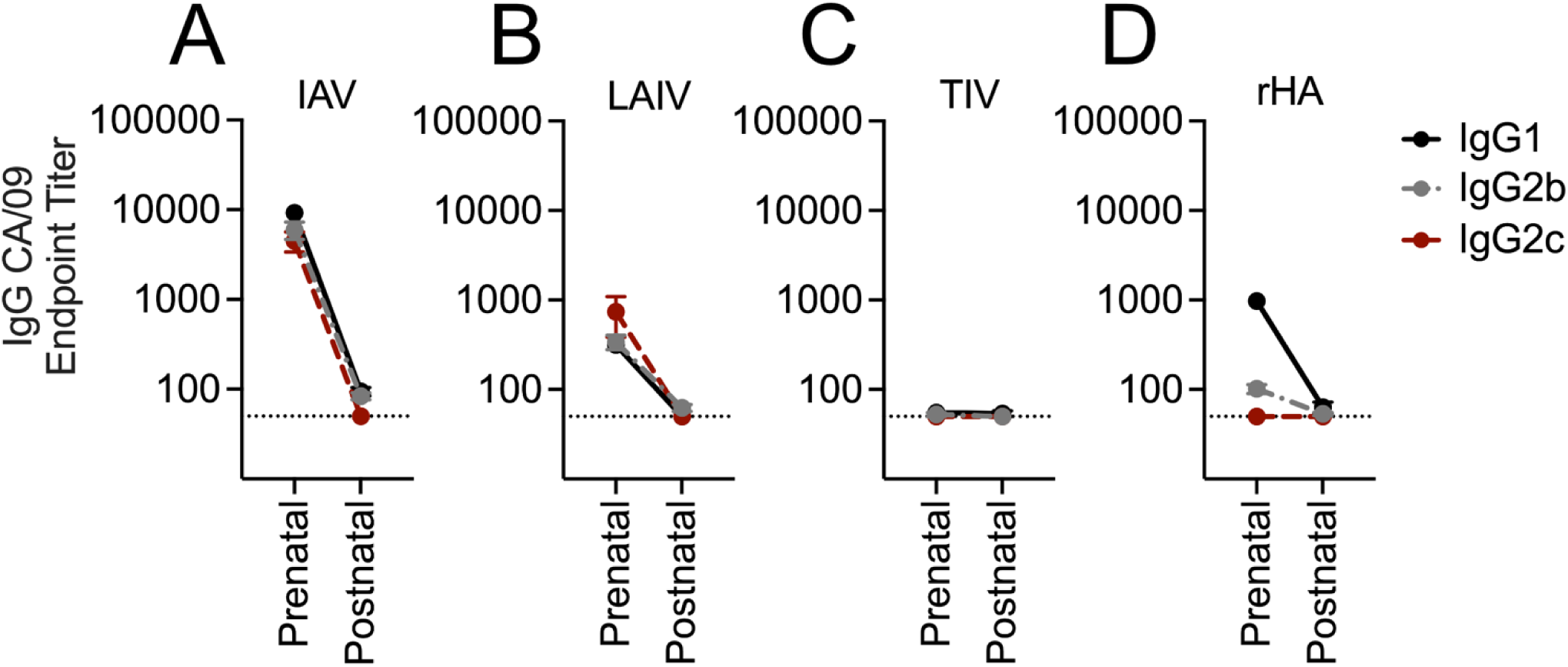
Maternal transfer of IgG2c viral-specific antibodies prenatally is associated with increased protection from lethal influenza virus infection. Antibody levels in sera were quantified via viral-specific ELISA using IgG1, IgG2b, and IgG2c-specific secondaries. Pre-challenge titers for cross-fostered prenatal or postnatal groups are shown for (A) IAV sub-lethal infection, (B) LAIV vaccination, (C) TIV vaccination, and (D) rHA vaccination with n=10 mice/group.

While offspring born to TIV- or rHA-vaccinated dams had other IgG subclass antibodies, these groups had no IgG2c viral-specific titers present in their serum (**Figures 6C-D, Supplementary Table 3**), which could begin to explain the diminished survival rates observed in offspring from these groups when not born to and nursed by the same dam (**Figure 3H**, **Figure 5F**). However, we observed high IgG1 viral-specific ELISA titers in the serum of pups born to rHA-vaccinated dams but breastfed by unvaccinated dams (prenatal transfer). This finding could account for the 20% increase in the probability of survival in this group as compared to TIV (**Figure 5F**, **Figures 6C-D, Supplementary Table 3**), suggesting a role for IgG1 subclass antibodies in mediating protection from challenge in our model. Overall, prenatal transfer was primarily enriched with IgG1 for all vaccination groups, except for pups born to TIV-vaccinated dams, yet the levels of IgG2c and IgG2b varied depending on the vaccination platform (**Figure 6, Supplementary Table 3**). Together, these data suggest that maternal exposure to distinct influenza vaccine platforms can elicit different antibody profiles that are preferentially transferred prenatally (**Figure 5**, **Figure 6**). Beyond influenza, these findings can apply to other respiratory viral infections and are vital to consider when seeking to improve the immunogenicity of current vaccine regimens during pregnancy to provide efficient and longitudinal protection in the offspring.

## Discussion

Given the difficulties in performing extensive vaccine studies with longitudinal sampling in pregnant women, mouse models of pregnancy have provided an opportunity to better understand the elicitation of humoral responses during pregnancy upon infection or vaccination.^36^ However, available preclinical studies still provide limited information on how additional influenza vaccine platforms may influence immunogenicity in dams and subsequently, passive immunity in their offspring. Here, we sought to investigate the efficacy and immunogenicity of distinct influenza vaccine platforms provided during pregnancy along with the kinetics and functionality of passively transferred matAbs. First, we monitored pregnant dams infected with IAV or vaccinated with TIV, LAIV, and rHA (**Supplementary Figure 1**). Upon vaccination with any of these vaccine platforms, we did not observe any adverse effects in weight or the presence of clinical scores in pregnant mice.^16,17,20,29^ However, we did observe infection-related weight loss in dams inoculated with a sub-lethal dose of A/CA/04/09 (**Supplementary Figure 1**). Additionally, the vaccine platforms used for these studies did not seem to alter the average weight of pups from each litter nor hinder growth in any of the groups when monitored until adulthood, or eight weeks of age (**Figure 4B**). These initial data further highlight an already established and vital finding that vaccination is safe and highly recommended during pregnancy.^15^

The effect of pregnancy on host immunity has long been appreciated as alterations during the gestational period, such as changes in hormones or the inflammatory secretome, can alter broad immune responses to infection, and immunization with a split-inactivated influenza vaccine.^36–39^ Our study extends this work and demonstrates variable outcomes depending on which influenza vaccine platform was used to immunize pregnant dams (**Figure 1**). While all vaccinated or previously infected groups exhibited improved protection from a lethal influenza challenge, LAIV and rHA vaccination along with IAV infection resulted in a significantly increased chance of survival (**Figure 1H**), a result that aligned with the quantity (**Figure 1B**) and quality of antibody levels in these groups (**Figure 1D**). Upon immunization of non-pregnant age-matched female mice with the same platforms, viral-specific IgG titers were higher for TIV-vaccinated mice as compared to dams immunized while pregnant which resulted in an 80% probability of survival in this group as compared to 20% in the latter (**Figures 1H**, **Figures 2B-H**). This differing but potentially interdependent contribution of pregnancy status upon immunization with a split-inactivated vaccine has been observed whereby B-cell frequency significantly decreases over time in women vaccinated during pregnancy.^40^ Yet, further study into the role of cellular immune responses is needed, especially in understanding whether there are additional impairments in B cell activation, proliferation, or differentiation due to the pregnancy or post-partum status.

Murine studies previously showed that vaccination against influenza during pregnancy enhances the transfer of viral-specific IgG and subsequent protection to offspring.^22,23^ Yet, most studies have only evaluated the role of matAbs when vaccinating the offspring at an early age.^21,22,27,41^ Even fewer studies have examined the application of additional immunization platforms other than an inactivated influenza virus vaccine or IAV infection.^22,27,28,42,43^ In our model, three-week-old offspring born to infected or LAIV-vaccinated dams exhibited high viral-specific IgG ELISA titers with neutralizing capacity leading to increased protection and lower pulmonary viral burden from subsequent IAV infection as compared to pups born to unvaccinated mice (**Figure 3**). Offspring born to IAV-infected, rHA- and LAIV-vaccinated dams had the highest transfer of viral-specific IgG antibodies from their mothers (**Figure 3B**) indicating that the vertically transferred matAbs elicited by these platforms play a role in the offspring’s protection against influenza. This finding is further strengthened by the observed reduction in lung viral titers at 6dpi (**Figure 3E**) and protection against infection with a lethal dose (10xMLD_50_) of influenza virus (**Figures 3F-3H**). However, offspring born to TIV-vaccinated dams exhibited suboptimal survival upon challenge (**Figure 3H**), which aligns with the absence of matAbs in this group before the challenge (**Figure 3B-3D**). Recently, work has illustrated how vertically transferred maternal immune cells may also play a role in protection against early-life infections.^44,45^ It would be advantageous to dissect what contribution, if any, these maternal microchimeric cells (MMc) may have in protecting the offspring in our model, and whether distinct influenza vaccine platforms or natural IAV infection harbor specific MMc profiles.

Previously, work by Chronopoulos et al. concluded that both prenatal and postnatal transfer of matAbs from IAV-infected dams to their 3-week-old offspring was required for protection against influenza.^42^ However, the offspring were not challenged with a live virus at this age. We found that infection or immunization with different influenza vaccines resulted in a preference towards prenatal matAb transfer to provide protective immunity in their offspring (**Figure 5**). While IgG is the primary antibody isotype transferred during pregnancy, not all IgG subclasses may be equally transferred. We also sought to identify the IgG subclass antibody profile present in the offspring’s serum before infection (**Figure 6, Supplementary Table 3**). Antibodies of the IgG2c subclass are known to stimulate or activate Fc receptor-mediated effector functions leading to improved viral clearance as compared to antibodies of additional IgG subclasses. Offspring with antibody profiles biased towards elevated IgG2c titers also had higher rates of survival (**Figure 5F-H, Supplementary Table 3**) suggesting a role for these antibodies in mediating protection from influenza possibly through increased antibody effector cell function as compared to other IgG subclasses. These findings aid in better understanding the maternal antibody profiles elicited via distinct influenza vaccines in a murine model and provide better context for the type of responses needed for protection in their offspring during early life.

While our studies successfully dissect the contribution of each influenza vaccine platform in dams and their offspring against influenza challenge or infection, there are limitations. First, it is vital to consider how inoculation dose may play a role in these outcomes. In our studies, we used a lethal infectious dose, or 10xMLD_50_, which may not reflect an accurate infection dose upon IAV infection in humans. It would be of interest to assess if different inoculation doses would alter any survival outcomes after vaccination and subsequent challenges. While we found distinct influenza vaccines can elicit unique antibody profiles when applied in our model, there are differences in the expression of Ig isotypes between mice and humans, and direct correlations between subtypes within classes in each species are difficult to make.^46^ If possible, it would be advantageous to assess the elicitation of distinct antibody responses upon vaccination with the available vaccines in a human cohort. The mice used in these studies were naïve and had no pre-existing immunity, so our findings isolate the elicitation of humoral responses to one or two immunizations or infections and the response to subsequent influenza challenges.

The relationship between pregnancy and infectious disease is not unique to the influenza virus. Epidemiological data and studies in pregnant mice demonstrate pregnancy is a risk factor for severe COVID-19, and vaccination has been associated with improved outcomes.^48,49^ In the search for a universal influenza vaccine, these findings highlight the functionality of maternal antibodies elicited via distinct vaccine delivery systems and are vital to refine vaccine development pipelines for this high-risk group. Our results are in alignment with recent advances such as Pfizer’s maternal respiratory syncytial virus (RSV) vaccine.^50^ Future studies must be performed to untangle cellular-mediated responses to each influenza vaccine platform in pregnant dams and their offspring, how they may influence influenza pathogenicity within the lung tissue to mediate protection, and other host behaviors that may impact the formation of primary and recall responses upon exposure to the antigen-of-interest using distinct vaccine delivery systems.

## Acknowledgments

The authors gratefully acknowledge the expert technical assistance and scientific discussions from lab members of the Schultz-Cherry laboratory at St. Jude Children’s Research Hospital. Thank you to the St. Jude Animal Resource Center and Center for In Vivo Imaging and Therapeutics for their helpful experimental assistance. This work was supported by ALSAC, the National Institute of Allergy and Infectious Diseases (NIAID) at the National Institutes of Health (NIH), by the NIAID Collaborative Influenza Vaccine Innovation Centers (CIVIC) contract 75N93019C00052 and contract 75N93021C00016 for the St. Jude Center of Excellence for Influenza Research and Response (CEIRR) to S. S.-C. E.K.R. was supported by NIAID T32AI106700-07. A.V.P. was supported by NIAID F31AI161986 and by the St Jude Graduate School for Biomedical Sciences. The content is solely the responsibility of the authors and does not necessarily represent the official views of the National Institutes of Health.

## Author contributions

Conceptualization – A.V.P., E.K.R.,S.S-C.; Methodology – A.V.P., E.K.R.,S.S-C.; Validation – A.V.P., E.K.R.,S.S-C.; Formal analysis– A.V.P., E.K.R.,S.S-C.; Investigation – A.V.P., E.K.R., S.C., B.L., T.B., L.L., B.S., T.C., P.H.B, K.T.W, M.J., V.M..; Resources – A.V.P., E.K.R., S.C., B.L., T.B., L.L.,B.S.,T.C.,P.H.B, K.T.W, M.J., V.M., S.S-C; Data curation – A.V.P., E.K.R.,S.D.C.; Writing - original draft – A.V.P., E.K.R.,S.D.C.; Writing – review and editing – All authors; Visualization – A.V.P., E.K.R.; Supervision – S.S-C.; Project administration – A.V.P., E.K.R., S.S-C.; Funding acquisition –S.S-C.. A.V.P., E.K.R. share first authorship and contributed equally to this work.

## Materials & Correspondence

Any requests for reagents, materials, or correspondence should be directed to the corresponding author at

## Materials and Methods

### Animal husbandry and ethics

All procedures were approved by the St. Jude Children’s Research Hospital Institutional Animal Care and Use Committee (IACUC) and complied with the Guide for the Care and Use of Laboratory Animals.

### Vaccinations

Timed-pregnant and non-pregnant 8-week-old C57BL/6J female mice were purchased from Jackson Laboratories (JAX 000664) and screened upon arrival via ultrasound to confirm pregnancies and gestational age. The vaccine schematic is shown in **Figure 1A**. Briefly, at embryonic day 8 (E8), mice were lightly anesthetized using 3% inhaled isoflurane and intramuscularly (IM) vaccinated with TIV (BEI NR-31063 at 3μg/HA/mouse) or 3μg of purified recombinant H1 (A/California/04/2009, SinoBiologicals 11055-V08B) and H3 (A/Hong Kong/4801/2014, SinoBiologicals 40555-V08B) HAs mixed with Addavax (Cat #ADX-40-02, Invitrogen) at a 1:1 ratio in the right hind leg via prime-boost with the boost occurring 3 weeks post-prime. Mice were also lightly anesthetized and intranasally (in) administered a sub-lethal dose of A/California/04/2009 (A/CA/04/09) virus at 50 plaque-forming units (pfu) or 10^6^ TCID_50_ of LAIV (cold-adapted-A/California/04/2009 in Ann Arbor backbone, CDC) in 25 µl PBS.^51^ Sera were collected, via retroorbital bleeds, at baseline, before birth (9 days post-vaccination or approximately at E17), and pre-challenge (5 weeks post-vaccination or 21 days postpartum).

### Challenge Studies

Pups were weaned from vaccinated or non-vaccinated dams at three weeks of age, housed by sex, and randomized for infection studies at three or eight weeks of age. All data reflect both male and female offspring. Non-vaccinated mice served as controls. After weaning pups at 3 weeks of age, dams were challenged with 10xMLD_50_ dose of A/CA/04/09 virus in 25 μl PBS for dams, 8-week-old male and female pups, and 3-week-old male pups. However, the virus was administered in 10 μl PBS for 3-week-old female pups due to significant size differences between both sexes. The inoculum was administered under light anesthesia (3% inhaled isoflurane). Weights were recorded every other day for 14 days. Clinical signs were scored daily as follows: 0=no observable signs, 1=active, squinting, hunching or scruffy appearance; 2=squinting and hunching, active only when stimulated; 3=excessive hunching, squinting, not active when stimulated; 4=hind-limb paralysis, shivering, rapid breathing, moribund. Mice losing more than 30% of body weight and reaching a clinical score of 3 were humanely euthanized. Tissues and sera were collected on days 3 and 6 post-challenge and processed immediately and stored at −80°C for future analysis. Mouse lethal dose-50 (MLD_50_) titers were determined in pregnant and postpartum dams, 3-week-old, and 8-week-old C57BL/6J male and female mice, respectively, to standardize inoculums. The number of mice per experiment is noted in the figure legends and text. Data is a summary of three independent experiments.

### Cross-Fostering

To determine whether protective passive immunization from vaccinated mothers occurred prenatally or postnatally, we performed cross-fostering and litter swap experiments as outlined in **Figure 5**, adapted from Chronopoulos et al.^42^ Briefly, after giving birth, unvaccinated and vaccinated dams were separated from their litters and placed in separate cages. Pups were carefully transferred using clean tweezers and rolled in the bedding of the new dam and placed in the same spot as the switched pups. Once litters were successfully transferred, dams were returned to their original cages and scruffed to promote urination and prevent rejection. After 3 weeks, pups were weaned, housed by sex, and infected as described above. All data reflect both male and female offspring. The number of mice per experiment is noted in the figure legends and text and data is a summary of two independent experiments.

### Viral Titer Determination

Viral titer was determined as previously reported.^52^ Briefly, Madin-Darby canine kidney (MDCK, ATCC CCL-34) cells were plated in 96-well tissue culture plates and incubated until confluent. After washing with PBS, cells were infected with 10-fold serial dilutions of whole lung homogenates (bead beat in 1mL 1X PBS) in 100 μL of Minimal essential media (MEM) supplemented with 0.75% BSA and 1 ug/ml TPCK-trypsin and incubated for 3 days at 37°C, 5% CO_2_. Cell supernatant was scored using hemagglutinin (HA) endpoint. Next, cells were fixed with 3.7% paraformaldehyde and stained with 1% crystal violet. HA endpoints were corroborated by the appearance of cytopathic effects. Tissue culture infectious dose-50 (TCID_50_) was determined by Reed and Muench method.^53^

### Antibody Titers

#### Whole virus IgG ELISA

Flat-Bottom Immuno Nonsterile 96-well Plates 4HBX (ThermoScientific, #3855) were coated with 50μL/well of 5μg/mL purified A/CA/04/09 diluted in 1X PBS and incubated overnight at 4°C. The next day, plates were washed 3x with 200μL of PBS containing 0.1% Tween-20 (PBS-T) and blocked with 200μLs of blocking solution (3% goat serum [Gibco, #16210072], 0.5% Non-Fat Milk Powder [AmericanBio, #AB10109] in PBS-T) for 1 hour at room temperature. After blocking, two-fold dilutions of sera were added to the wells and incubated for 2 hours at room temperature. Plates were then washed 3x with 200μLs of PBS-T per well and anti-mouse IgG HRP-conjugated secondary (Sigma, #AP308P) diluted at 1:3000 in blocking solution was added to each well and incubated for 1 hour at room temperature. For IgG subbclass specific ELISAs, anti-IgG1 (Abcam #AB97240, 1:5000 dilution), IgG2b (Abcam #AB97250, 1:5000 dilution), and IgG2c (Southern Biotech #1079-05,1:3000 dilution) were used. After washing, Sigma Fast OPD (Millipore Sigma, #P9187) was added to each well, incubated for 10 minutes, and stopped with 3M HCl. Absorbance at 490 nm was read on a spectrophotometer and analyzed using Microsoft Excel. Data are shown as endpoint titers, representing the highest dilution at which a signal was still captured. Samples were run in duplicate.

#### Hemagglutination inhibition assay (HAI)

HAI assays were performed as described.^54^ Briefly, sera were treated overnight with receptor-destroying enzyme (Denka Seiken, Co., Ltd.) and 2-fold serially diluted in 25 µL of PBS in duplicate in a 96-well v-bottom plate (Corning, #3897). A/CA/04/09 virus was adjusted to 4 hemagglutination units (HAU) and 25 µl were added to the sera dilutions and incubated for 30 min at 4°C. Finally, 50 µl of 0.5% turkey red blood cells (Rockland, #R408-0050) were added to each of the wells and the plate was stored at 4°C for 30 min. after which the assay was read. Samples were run in duplicate.

#### Luminescent Microneutralization Assay (MN)

Luminescent MNs were performed as described.^55^ Briefly, MDCK cells were plated at 3×10^4^ cells/well in white Bottom, 96-well tissue culture plates (Costar, Cat #3917) and incubated at 37°C, 5% CO_2_ overnight. Two-fold serial dilutions of RDE-treated sera in microneutralization media (50µLs, MEM with 1% L-glutamine, 1x Pen/Strep, and 1% BSA) were incubated with 2 × 10^3^ TCID_50_/NanoLuc A/California/04/2009 (CA/09-PA NLuc) virus for 1 hour at 37°C, 5% CO_2_ before being added to pre-washed MDCK cells. Cells were incubated at 37°C, 5% CO_2_ for 18 hours, followed by removal of the media, and a freeze-thaw at −80°C for at least 30 minutes to lyse the cells. NanoGlo (Promega, #N1150) reagent was added at 25µLs per well) and plates were read on a luminometer with a gain setting of 100 to prevent variability between plates. All samples were run in duplicate. The neutralization titer for each serum sample was expressed as the reciprocal of the highest dilution at which virus infection was blocked as the formula below:

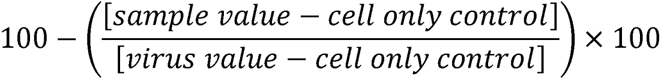

#### Statistical analyses

Power analysis was performed to ensure study rigor with β=0.2. Data were compiled and organized using Microsoft Excel and GraphPad Prism v9.3.1. Statistical analysis was performed using GraphPad Prism v9.3.1 as described in the figure legends. Statistical significance was defined as α=0.05 and, unless otherwise specified, is comparing the unvaccinated dams or pups born to unvaccinated dams to the other experimental groups.

## References

1. Woolston, W. J. & Conley, D. O. Epidemic Pneumonia (Spanish Influenza) in Pregnancy: Effect in One Hundred and One Cases. J. Am. Med. Assoc. 71, 1898–1899 (1918).

2. Sappenfield, E., Jamieson, D. J. & Kourtis, A. P. Pregnancy and susceptibility to infectious diseases. Infect Obstet Gynecol 2013, 752852 (2013).

3. Greenberg, M., Jacobziner, H., Pakter, J. & Weisl, B. A. Maternal mortality in the epidemic of Asian influenza, New York City, 1957. Am J Obstet Gynecol 76, 897–902 (1958).

4. Rasmussen, S. A., Jamieson, D. J. & Bresee, J. S. Pandemic influenza and pregnant women. Emerg Infect Dis 14, 95–100 (2008).

5. Lieberman, R. W., Bagdasarian, N., Thomas, D. & Van De Ven, C. Seasonal influenza A (H1N1) infection in early pregnancy and second trimester fetal demise. Emerg Infect Dis 17, 107–9 (2011).

6. Prasad, N. et al. Influenza-Associated Outcomes Among Pregnant, Postpartum, and Nonpregnant Women of Reproductive Age. J Infect Dis 219, 1893–1903 (2019).

7. Crow, T. J. & Done, D. J. Prenatal exposure to influenza does not cause schizophrenia. Br J Psychiatry 161, 390–3 (1992).

8. Brown, A. S. Epidemiologic studies of exposure to prenatal infection and risk of schizophrenia and autism. Dev Neurobiol 72, 1272–6 (2012).

9. Ellman, L. M., Yolken, R. H., Buka, S. L., Torrey, E. F. & Cannon, T. D. Cognitive functioning prior to the onset of psychosis: the role of fetal exposure to serologically determined influenza infection. Biol Psychiatry 65, 1040–7 (2009).

10. Brown, A. S. et al. Serologic evidence of prenatal influenza in the etiology of schizophrenia. Arch Gen Psychiatry 61, 774–80 (2004).

11. Haberg, S. E. et al. Risk of fetal death after pandemic influenza virus infection or vaccination. N Engl J Med 368, 333–40 (2013).

12. Laake, I. et al. Risk of pregnancy complications and adverse birth outcomes after maternal A(H1N1)pdm09 influenza: a Norwegian population-based cohort study. BMC Infect Dis 18, 525 (2018).

13. Le, T. V. et al. Fatal avian influenza A(H5N1) infection in a 36-week pregnant woman survived by her newborn in Soc Trang Province, Vietnam, 2012. Influenza Respir Viruses 13, 292–297 (2019).

14. Honce, R., Wohlgemuth, N., Meliopoulos, V. A., Short, K. R. & Schultz-Cherry, S. Influenza in High-Risk Hosts-Lessons Learned from Animal Models. Cold Spring Harb Perspect Med 10, (2020).

15. Vaccines against influenza WHO position paper - November 2012. Wkly Epidemiol Rec 87, 461–76 (2012).

16. Zaman, K. et al. Effectiveness of maternal influenza immunization in mothers and infants. N Engl J Med 359, 1555–64 (2008).

17. Nunes, M. C. et al. Duration of Infant Protection Against Influenza Illness Conferred by Maternal Immunization: Secondary Analysis of a Randomized Clinical Trial. JAMA Pediatr 170, 840–7 (2016).

18. Leuridan, E. & Van Damme, P. Passive transmission and persistence of naturally acquired or vaccine-induced maternal antibodies against measles in newborns. Vaccine 25, 6296–304 (2007).

19. Pou, C. et al. The repertoire of maternal anti-viral antibodies in human newborns. Nat Med 25, 591–596 (2019).

20. Albrecht, M. & Arck, P. C. Vertically Transferred Immunity in Neonates: Mothers, Mechanisms and Mediators. Front Immunol 11, 555 (2020).

21. Hwang, S. D. et al. Protection of pregnant mice, fetuses and neonates from lethality of H5N1 influenza viruses by maternal vaccination. Vaccine 28, 2957–64 (2010).

22. van der Lubbe, J. E. M. et al. Maternal antibodies protect offspring from severe influenza infection and do not lead to detectable interference with subsequent offspring immunization. Virol J 14, 123 (2017).

23. Esser, E. S. et al. Microneedle patch delivery of influenza vaccine during pregnancy enhances maternal immune responses promoting survival and long-lasting passive immunity to offspring. Sci Rep 7, 5705 (2017).

24. Beagley, K. W. & Gockel, C. M. Regulation of innate and adaptive immunity by the female sex hormones oestradiol and progesterone. FEMS Immunol. Med. Microbiol. 38, 13–22 (2003).

25. Jamieson, D., Theiler, R. & Rasmussen, S. Emerging Infections and Pregnancy. Emerg. Infect. Dis. 12, 1638–1643 (2006).

26. Kim, J. C. et al. Severe pathogenesis of influenza B virus in pregnant mice. Virology 448, 74–81 (2014).

27. Creisher, P. S. et al. Influenza subtype-specific maternal antibodies protect offspring against infection but inhibit vaccine-induced immunity and protection in mice. Vaccine 40, 6818–6829 (2022).

28. Suguitan, A. L. et al. Influenza H1N1pdm-specific maternal antibodies offer limited protection against wild-type virus replication and influence influenza vaccination in ferrets. Influenza Respir Viruses 8, 169–76 (2014).

29. Toback, S. L. et al. Maternal outcomes among pregnant women receiving live attenuated influenza vaccine: Outcomes in women who received LAIV while pregnant. Influenza Other Respir. Viruses 6, 44–51 (2012).

30. Huber, V. C. et al. Distinct contributions of vaccine-induced immunoglobulin G1 (IgG1) and IgG2a antibodies to protective immunity against influenza. Clin. Vaccine Immunol. CVI 13, 981–990 (2006).

31. Fink, A. L., Engle, K., Ursin, R. L., Tang, W.-Y. & Klein, S. L. Biological sex affects vaccine efficacy and protection against influenza in mice. Proc. Natl. Acad. Sci. U. S. A. 115, 12477–12482 (2018).

32. Gerhard, W., Mozdzanowska, K., Furchner, M., Washko, G. & Maiese, K. Role of the B-cell response in recovery of mice from primary influenza virus infection. Immunol. Rev. 159, 95–103 (1997).

33. Mozdzanowska, K., Furchner, M., Washko, G., Mozdzanowski, J. & Gerhard, W. A pulmonary influenza virus infection in SCID mice can be cured by treatment with hemagglutinin-specific antibodies that display very low virus-neutralizing activity in vitro. J. Virol. 71, 4347–4355 (1997).

34. Coutelier, J. P., van der Logt, J. T., Heessen, F. W., Vink, A. & van Snick, J. Virally induced modulation of murine IgG antibody subclasses. J. Exp. Med. 168, 2373–2378 (1988).

35. Coutelier, J. P., van der Logt, J. T., Heessen, F. W., Warnier, G. & Van Snick, J. IgG2a restriction of murine antibodies elicited by viral infections. J. Exp. Med. 165, 64–69 (1987).

36. Vazquez-Pagan, A. & Schultz-Cherry, S. Serological Responses to Influenza Vaccination during Pregnancy. Microorganisms 9, 2305 (2021).

37. Engels, G. et al. Pregnancy-Related Immune Adaptation Promotes the Emergence of Highly Virulent H1N1 Influenza Virus Strains in Allogenically Pregnant Mice. Cell Host Microbe 21, 321–333 (2017).

38. Aghaeepour, N., et al. An immune clock of human pregnancy. Sci Immunol 2, (2017).

39. Vazquez-Pagan, A., Honce, R. & Schultz-Cherry, S. Impact of influenza virus during pregnancy: from disease severity to vaccine efficacy. Future Virol. 15, 441–453 (2020).

40. Kay, A. W. et al. Pregnancy Does Not Attenuate the Antibody or Plasmablast Response to Inactivated Influenza Vaccine. J. Infect. Dis. 212, 861–870 (2015).

41. Sweet, C., Jakeman, K. J. & Smith, H. Role of milk-derived IgG in passive maternal protection of neonatal ferrets against influenza. J Gen Virol 68 (Pt 10), 2681–6 (1987).

42. Chronopoulos, J., Martin, J. G. & Divangahi, M. Transplacental and Breast Milk Transfer of IgG1 Are Both Required for Prolonged Protection of Offspring Against Influenza A Infection. Front. Immunol. 13, 823207 (2022).

43. Paquette, S. G. et al. Influenza Transmission in the Mother-Infant Dyad Leads to Severe Disease, Mammary Gland Infection, and Pathogenesis by Regulating Host Responses. PLoS Pathog 11, e1005173 (2015).

44. Stelzer, I. A., Thiele, K. & Solano, M. E. Maternal microchimerism: lessons learned from murine models. J Reprod Immunol 108, 12–25 (2015).

45. Stelzer, I. A. et al. Vertically transferred maternal immune cells promote neonatal immunity against early life infections. Nat Commun 12, 4706 (2021).

46. Mestas, J. & Hughes, C. C. W. Of mice and not men: differences between mouse and human immunology. J. Immunol. Baltim. Md 1950 172, 2731–2738 (2004).

47. Ander, S. E., Diamond, M. S. & Coyne, C. B. Immune responses at the maternal-fetal interface. Sci Immunol 4, (2019).

48. Zhu, G. et al. Delayed Antiviral Immune Responses in Severe Acute Respiratory Syndrome Coronavirus Infected Pregnant Mice. Front. Microbiol. 12, 806902 (2022).

49. Villar, J. et al. Pregnancy outcomes and vaccine effectiveness during the period of omicron as the variant of concern, INTERCOVID-2022: a multinational, observational study. The Lancet S0140673622024679 (2023) doi:10.1016/S0140-6736(22)02467-9.

50. Simões, E. A. F. et al. Prefusion F Protein–Based Respiratory Syncytial Virus Immunization in Pregnancy. N. Engl. J. Med. 386, 1615–1626 (2022).

51. Chen, Z. et al. Generation of live attenuated novel influenza virus A/California/7/09 (H1N1) vaccines with high yield in embryonated chicken eggs. J. Virol. 84, 44–51 (2010).

52. Cline, T. D., Karlsson, E. A., Seufzer, B. J. & Schultz-Cherry, S. The hemagglutinin protein of highly pathogenic H5N1 influenza viruses overcomes an early block in the replication cycle to promote productive replication in macrophages. J. Virol. 87, 1411–1419 (2013).

53. Reed, L. J. & Muench, H. A SIMPLE METHOD OF ESTIMATING FIFTY PER CENT ENDPOINTS12. Am. J. Epidemiol. 27, 493–497 (1938).

54. Karlsson, E. A. et al. Obesity Outweighs Protection Conferred by Adjuvanted Influenza Vaccination. Mbio 7, (2016).

55. Karlsson, E. A. et al. Visualizing real-time influenza virus infection, transmission and protection in ferrets. Nat. Commun. 6, 6378 (2015).

