## Supplemental figures and information for "Maternal immunization with distinct influenza vaccine platforms elicits unique antibody profiles that impact the protection of offspring"

### **Supplementary material**

**Supplementary Figures and Legends**

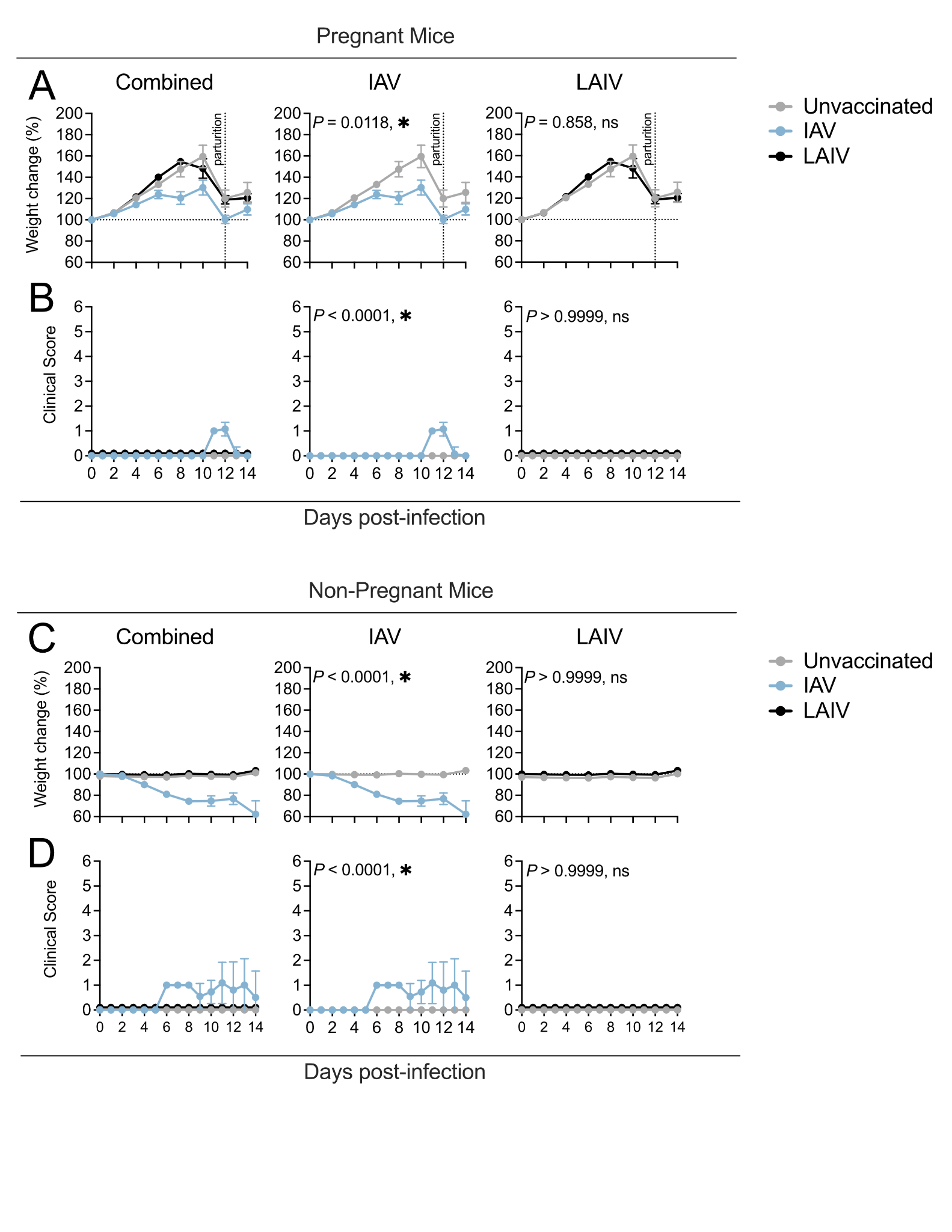

**Supplementary Figure 1. No adverse maternal outcomes were associated with LAIV vaccination in pregnant mice.** (A-D) Morbidity in pregnant and non-pregnant vaccinated or infected mice compared to unvaccinated mice with n=15 mice/group. (A, C) Weight curves post-challenge with statistical comparisons made via mixed-effects model. Grey dotted line indicates a 0% deviation from weight at baseline. (B, D) Clinical scores were monitored for 14 days in pregnant (A, B) and non-pregnant mice (C, D) after intranasal administration of IAV or LAIV compared to unvaccinated mice with statistical comparisons made via a two-way ANOVA. Data are representative of three independent experiments and are graphed as means ± standard error. Statistical comparison tests are between unvaccinated and vaccinated/infected groups (n=15 mice/group) in (A-D).

**Supplementary Table 1. Summary of results upon maternal immunization and lethal influenza H1N1 virus challenge in post-partum dams.**

| Platform for maternal inoculation or immunization | Antibody Quantification^a^ | | | Lung Viral Load  (log_10_ TCID_50_/mL)^b^ | | Survival Rate (%) |
| --- | --- | --- | --- | --- | --- | --- |
|  | IgG CA/09 Endpoint Titer | Reciprocal HAI CA/09 Titer (1:X) | Reciprocal MN CA/09 Titer (1:X) | 3dpc^c^ | 6dpc |  |
| IAV | 47360 | 160 | 1920 | 1.0 ± 0 | 1.0 ± 0 | 100 |
| LAIV | 9600 | 10 | 320 | 5.248 ± 0.939 | 1.0 ± 0 | 100 |
| TIV | 271.43 | 10 | 40 | 5.437 ± 1.689 | 6.583 ± 0.824 | 20 |
| rHA | 39680 | 40 | 5120 | 1.0 ± 0 | 2 ± 0.707 | 100 |

^a^Data illustrated corresponds to the timepoint before challenge (or pre-challenge) and represents the mean values for each group.

^b^Results are expressed as the mean ± SD of tested mice in each group.

^c^dpc=days post-infection/days post-challenge

**Supplementary Table 2. Summary of results in offspring born to immunized or inoculated dams upon lethal influenza H1N1 virus challenge.**

| Platform for maternal inoculation or immunization | Antibody Quantification^a^ | | | Lung Viral Load  (log_10_ TCID_50_/mL)^b^ | | Survival  Rate (%) | | Timing of Maternal Antibody Transfer^c^ | | | |
| --- | --- | --- | --- | --- | --- | --- | --- | --- | --- | --- | --- |
|  |  |  |  |  |  |  |  | Prenatal | | Postnatal | |
|  | IgG CA/09 Endpoint Titer | Reciprocal HAI CA/09 Titer (1:X) | Reciprocal MN CA/09 Titer (1:X) | 3dpi^d^ | 6dpi | |  | IgG CA/09 Endpoint Titer^a^ | Survival Rate (%) | IgG CA/09 Endpoint Titer^a^ | Survival Rate (%) |
| IAV | 12800 | 160 | 240 | 4.038 ±1.468 | 2.0 ±1.414 | | 100 | 3460 | 100 | 97.2 | 40 |
| LAIV | 2160 | 10 | 120 | 4.118 ±1.837 | 6.416 ±0.276 | | 100 | 340 | 70 | 285 | 10 |
| TIV | 62.5 | 10 | 40 | 7 ±0.316 | 4.61 ±0.387 | | 20 | 55 | 30.7 | 56.5 | 31.25 |
| rHA | 3920 | 40 | 5120 | 7.125 ±0.735 | 6.4 ±0.561 | | 100 | 192.857 | 53.3 | 55 | 53.3 |

^a^Data illustrated corresponds to the timepoint before challenge (or pre-challenge) and represents the mean values for each group.

^b^Results are expressed as mean ± SD of tested mice in each group.

^c^Separation of data between prenatal and postnatal is concordant with the group separation as illustrated in Figure 5.

^d^dpi=days post-infection

**Supplementary Table 3. Summary of antibody subclass profiling in offspring born to immunized or inoculated dams.**

| Platform for maternal inoculation or immunization | Timing of Maternal Antibody Transfer^a^ | | | | | | | | |
| --- | --- | --- | --- | --- | --- | --- | --- | --- | --- |
|  | Prenatal | | | | | Postnatal | | | |
|  | IgG1 CA/09  Endpoint Titer^b^ | IgG2b CA/09  Endpoint Titer^b^ | IgG2c CA/09  Endpoint Titer^b^ | Survival Rate (%) | IgG1 CA/09  Endpoint Titer^b^ | | IgG2b CA/09  Endpoint Titer^b^ | IgG2c CA/09  Endpoint Titer^b^ | Survival Rate (%) |
| IAV | 9200 | 5960 | 4500 | 100 | 94.444 | | 83.333 | 50 | 40 |
| LAIV | 315 | 340 | 735.71 | 70 | 50 | | 62.5 | 50 | 10 |
| TIV | 55 | 52.5 | 50 | 30.7 | 53.846 | | 50 | 50 | 31.25 |
| rHA | 973.33 | 101.67 | 50 | 53.3 | 63.333 | | 53.333 | 50 | 53.3 |

^a^Separation of data between prenatal and postnatal is concordant with the group separation as illustrated in Figure 5.

^b^Data illustrated corresponds to the timepoint before challenge (or pre-challenge) and represents the mean values for each group.
